# Conformation Driven Enhancement of Neurolysin Activity in Presence of a Small Molecule Activator

**DOI:** 10.64898/2026.01.05.697776

**Authors:** Samik Bose, Asmaa Aly, Vardan T. Karamyan, Benjamin J. Orlando, Alex Dickson

**Affiliations:** Department of Computational Mathematics Science and Engineering, Michigan State University, East Lansing, MI 48824; Foundational Medical Studies, William Beaumont School of Medicine, Oakland University, Rochester, MI 48309; Department of Biochemistry and Molecular Biology, Michigan State University, East Lansing, Michigan 48824, USA

## Abstract

Neurolysin (Nln) is an M3 metallopeptidase that regulates neuropeptide concentration in the central nervous system. It has emerged as a therapeutic target for mitigating post-ischemic injury by hydrolyzing and inactivating several neuropeptides. It has been recently shown that small molecule activators, such as pyridine piperazine (Py-Pip) derivatives, can enhance Nln catalytic activity, facilitating hydrolysis of Nln substrate peptides. However, binding sites of these molecules and the mechanism of action remain unclear due to the dynamic nature of Nln. Here, we use molecular dynamics (MD) simulation of the apo and activator-bound Nln systems along with Markov state modeling to identify three binding sites of Py-Pip, and quantify their effects on the Nln conformational landscape and kinetics. Two of these sites have opposing functional outcomes: Binding inside the interhelical channel favors *closed* conformations by slowing down channel opening, thus allowing the substrates to orient favorably for the catalytic reaction, while binding at the exterior region of the channel stabilizes *open* states, potentially facilitating both substrate entry and cleaved product release. These findings suggest a possibility of a multi-site allosteric activation model in which distinct binding locations selectively modulate different steps within a catalytic cycle. This framework provides a structural and kinetic model for the action of small molecule Nln activators, offering insight into the design of cerebro-protective therapeutics that reduce neuropeptide accumulation after ischemic injury.

## Introduction

Neurolysin (Nln) is a zinc-metallopeptidase of the M3 family that hydrolyzes neuropeptides such as neurotensin (NT), substance P (SP) and bradykinin (BK) into smaller inactive peptides.^1–6^ Nln is abundantly present as monomers in cytosol, mitochondria and plasma membrane of the cells in the central nervous system. ^7,8^ The catalytic site of Nln is formed by a HIS-GLU-X-X-HIS motif, characteristic to various zinc dependent metallopeptidases, and a GLU residue from an opposing helix, participating in the coordination of the central zinc ion.^6^ The positively charged zinc ion strongly induces the GLU residues in the catalytic site, which in turn polarizes water molecules in the vicinity creating a favorable environment for water-mediated hydrolysis of the peptide bond. ^6,9^ The active site motif, shown in Fig. 1A, is conserved across the M3 class of metallopeptidase such as thimet oligopeptidase (TOP), angiotensin converting enzyme (ACE) and ACE2.^10,11^

**Figure 1:**
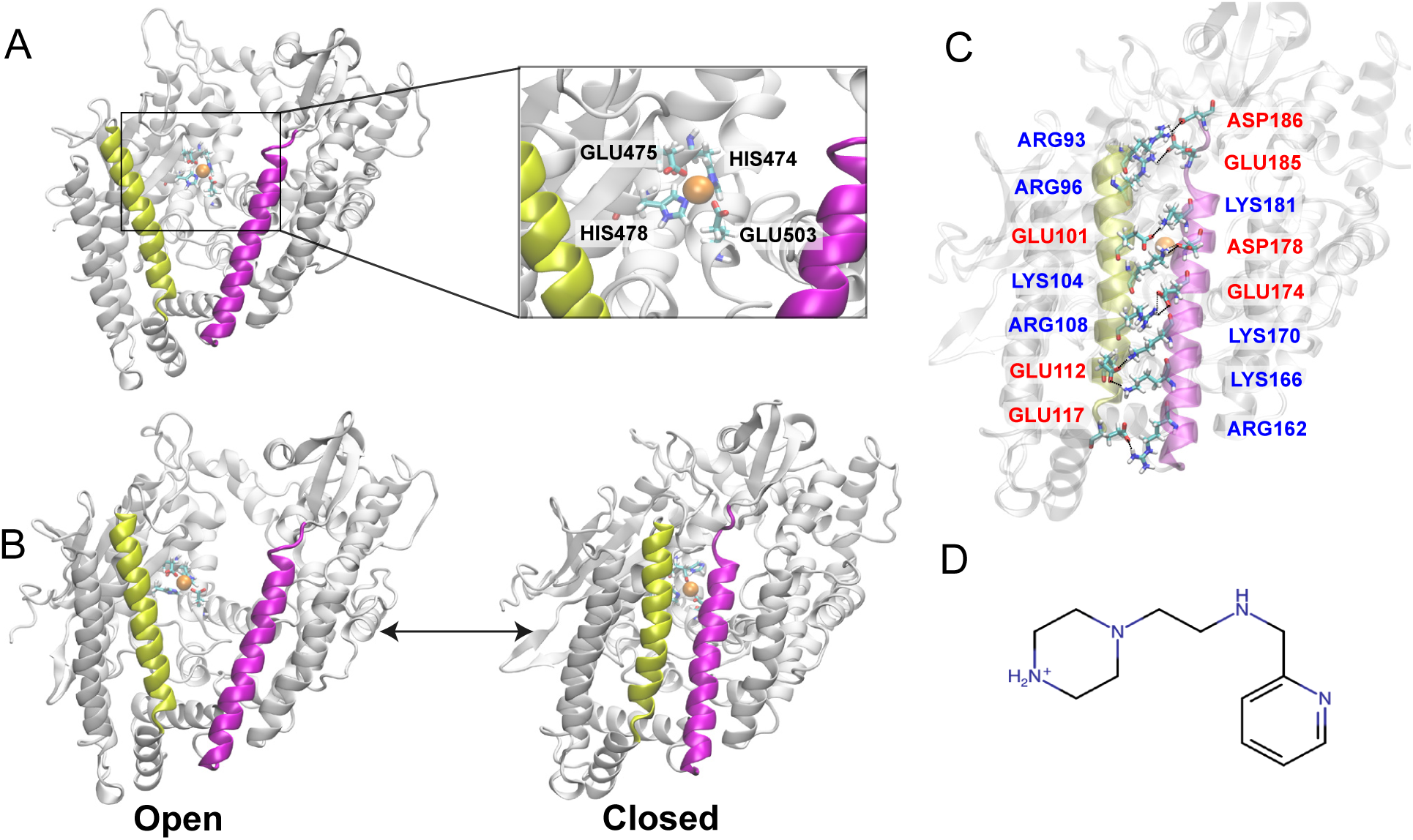
The structural detail of Nln and its activator Py-Pip molecule: (A) The catalytic motif of Nln (HEXXH…E) along with the zinc co-factor, (B) *Open* and *Closed* conformational states of Nln with the interhelical domains (*α* helices) highlighted in yellow and magenta, (C) The inter-domain salt-bridges stabilizing the closed conformation in the apo Nln with H-bond donors shown in blue and acceptors in red, (D) The chemical structure of Py-Pip molecule at 7.2 pH.

Nln has a deep channel running across the length of the molecule that divides the protein into two domains. The Nln active site cavity and its optimal structural features ensure the selectivity of the substrate polypeptides, which does not generally go beyond 17 amino acids in length.^6^ Nln and related metallopeptidases have been shown to possess inherent flexibility, with high-resolution structures of the enzyme being determined in two distinct *open* and *closed* conformational states as shown in Fig. 1B. In the *closed* state a number of salt-bridges are observed along the length of alpha-helices, forming a gate to the catalytic site (Fig. 1C). In a more dynamic *open* state, the two alpha-helix domains are significantly distant from each other increasing the volume of the active site cavity. Thus the open conformation of Nln favors substrate entry and the release of cleaved product, whereas the *closed* state tightly orients the substrate and the active site residues favoring the catalytic reaction. Such conformational switching between the *open* and *closed* states is important for several metallopeptidases to carry out their functions.^9,12,13^

The substrate neuropeptides of Nln are short chain polypeptides that modulate a variety of physiological processes, ranging from pain perception to the brain’s responses to stress and injury.^14–21^ Among them, NT, SP and BK have well-documented roles in the pathogenesis of stroke.^22^ Particularly, the extracellular concentrations of these neuropeptides are elevated and contribute to ischemia-induced brain injury.^1,3,23^ Notably, *in-vivo* over-expression of Nln in the mouse brain has been shown to reduce ischemic brain injury via increased clearance of these neuropeptides.^22^ On the other hand, inhibition of brain Nln after stroke leads to accumulation of these neuropeptides, aggravating ischemic brain injury.^22,24^ Based on these observations, Nln is deemed to be a therapeutic target for ischemic stroke^1,8,22^ and efforts are underway to develop drug-like small molecules that can enhance catalytic activity of this peptidase.^25–29^

These drug-like molecules are broadly categorized as Nln activators and should ideally follow the properties of standard small molecule drugs targeting the brain i.e., be stable upon peripheral administration, cross the blood-brain barrier (BBB) effectively, have high potency, are easy to manufacture, activate Nln without affecting the function of other pharmacological targets. Jayaraman and coworkers were first to discover His-His and His-Tyr dipeptides as hit molecules that enhance the catalytic activity of Nln against various peptide substrates. ^25^

They virtually screened the NCI Developmental Therapeutics Program Library against the rat Nln crystal structure (PDB ID: 1i1i) and analyzed the top 40 hits in an *in vitro* Nln activation assay that identified the two dipeptides. Based on this, Rahman and co-workers conduced detailed structure-activity relationship studies to identify the pharmacophore(s) in His-dipeptides and develop higher potency peptidomimetic analogs with improved drug-like properties.^26–28,30^ The presence of amide bonds in these compounds affects the lipophilicity and lowers the permeability through the blood-brain barrier.^31,32^ This was later overcome by introducing imidazole bioisostere compounds while preserving the HIS subunit from the dipeptides, which showed comparatively higher potency and blood-brain barrier penetration in structure-activity relationship studies.^30^ Key progress in Nln activator design was marked by the discovery of non-peptidomimetic pyridine-piperazine (Py-Pip) scaffold based compounds (Fig. 1D) which have sub-micromolar potency and increased brain penetration.^33,34^ The Py-Pip scaffold based molecules are shown to activate both recombinant human and rat Nln and in vitro favorable pharmacokinetic profiling of these molecules encourages their use as lead compounds for further *in-vivo* trials.^34^

While both chemotypes (peptidomimetic and Py-Pip analogs) offer promise in developing high-potency, brain permeable Nln activators with optimal drug-like properties, the underlying mechanism(s) by which these activators enhance the catalytic activity of Nln are not known. Plausible mechanisms for activation include: (i) inducing more open conformations that may help substrates to bind and/or products to release, ^9^ (ii) inducing more closed conformations that may help substrates to pack tightly in the catalytic site for the reaction,^6,13^ (iii) directly interacting with Nln residues near the catalytic site in a manner that promotes substrate ordering and reaction efficiency. Despite years of effort, there is currently no conclusive evidence defining the binding site(s) or mechanism of action for the Nln activators. Previous drug affinity responsive target stability (DARTS) and differential scanning fluorimetry (DSF) studies with dipeptide activators indicate the activators interact with the enzyme itself and not with the substrates.^25,26^ Such findings also align with the previously observed allosteric inhibition mechanism of Nln by a small molecule.^35–37^ Thus far, structural efforts using X-ray crystallography have failed to identify a binding site for the allosteric activators. Revealing the potential binding site(s) and understanding the mechanism of action of Nln activators are essential steps to design Nln activators as improved drug candidates. Such information will guide rational design of molecules with higher binding affinity while also preserving pharmacokinetic properties. Moreover, identification of binding site(s) is the first step towards understanding effect of the activators on Nln dynamics and overall catalytic potentiation.

While virtual screening and docking have previously been used to isolate hit compounds for Nln, these tools do not consider the effects of protein and water dynamics towards the binding of compounds, which is central to metallopeptidase activity. ^38^ Molecular dynamics (MD) simulation, on the other hand, is a powerful tool to study the conformational variation in metallopeptidases under different conditions, both in the apo enzyme and in the presence of a modulator.^36,39^ Furthermore, long and unbiased MD simulations can provide insight about the probable activator binding locations on the Nln. Since the relevance of the *open* and *closed* states of Nln are already well established in relation to enzymatic function, understanding the differential stability of the ensembles corresponding to these states in presence of the activator are crucial to decode the activator mechanism of action. In this context, a comparison between the mean first passage time (MFPT) of *open* ↔ *closed* inter-conversion in apo and Py-Pip bound Nln, along with the free energy landscape will provide the thermodynamic and kinetic insights of the activator induced conformational changes in Nln.

Here, we utilized MD simulations to understand the probable binding regions of the Py-Pip activator on Nln along with providing a plausible mechanism of action. To confirm the direct binding of Py-Pip in Nln and to assess the effect of binding towards the thermodynamic stability of the protein, we carry out differential scanning fluorimetry (DSF), a fluorescence-based thermal shift assay that reports ligand-induced changes in protein melting temperature. We construct the free energy landscape of Nln conformation space under apo and different activator-bound conditions in order to compare the relative stability of *open* and *closed* states. Additionally, we characterize the MFPT of the interconversion between *open* and *closed* states in presence of Py-Pip for understanding the kinetic relevance of the binding sites. By connecting activator binding with Nln conformational dynamics, our work provides mechanistic insights that may guide the design of next-generation cerebroprotective drugs.

## Methods

### Details of MD simulations

In this work, we have simulated six Nln systems: (i) apo Nln starting at the crystal structure, (ii) apo Nln starting from an *open* state, (iii) Nln with multiple copies of Py-Pip, (iv) Nln with single copy of Py-Pip bound at each of the three predicted binding sites. We have utilized the rat Nln crystal structure as the reference (PDB ID: 1i1i) for all of our simulations since the specific activator that we are examining here has been reported to show its activity on both recombinant human and rat Nln. Moreover, the rat Nln (PDB ID: 1i1i) has previously been used as a reference starting point for computational studies exploring inhibitor effects in Nln conformation.^36^ We set up all of our MD simulations using CHARMM-GUI^40^ to be run in the MD engine OpenMM-8.1^41^ with CHARMM36 force field.^42^ The force field parameters of Py-Pip molecule are generated using CGenFF tool with CHARMM36.^43^ The system size for all the simulations is a 90 Å cubic box with periodic boundary images that encompass the protein completely with at least 10 Å distance from the box edge to any point of the protein. All the systems are simulated in water with 0.15 M of added NaCl. Also, it is ensured that the charge of each system is neutralized with additional Na or Cl ions as per the requirement. For the Nln simulations in presence of Py-Pip activators, we have added 10 activator molecules randomly inside the simulation box, using the multi-component assembler utility of CHARMM-GUI.^44^ This system is denoted as Nln:10Py-Pip simulation henceforth. Routine energy minimization followed by NVT equilibration was performed for all the systems mentioned earlier in OpenMM to prepare them for production run. The NVT equilibrations were run for 250 ps at 300 K temperature with 1 fs time step. Further NPT equilibrations were run for 10 ns at 300 K and 1 atm pressure governed by a Monte Carlo barostat with 2 fs time step, using a stochastic Langevin integrator. The independent production runs for each system were continued at the same temperature and pressure. Details of the number and duration of the MD replicates for each system are given in SI Table S1. Total simulation times range from 12.5 to 18 *µ*s for each of the six systems, using 10 to 20 replicates. For all systems, equally spaced snapshots at 1 ns were extracted from simulation trajectories to carry out further analysis.

### Markov State Modeling with MD ensembles

A Markov state model (MSM) describes the evolution of a system over a period of time, known as lag time, in terms of transitions between discrete states and identifies slower modes in an ensemble. We have used MSMs to obtain mean first-passage times (MFPTs) of interconversion between the *open* and *closed* ensembles in apo Nln and different Py-Pip bound Nln systems. Direct estimation of MFPTs from MD trajectories is often noisy for these slow and sparse transitions. MSM utilizes short-time transition counts across the states to yield robust kinetic estimates of the same underlying processes. For MD simulations, a common approach is to use states that represent sets of conformations obtained by clustering the atomic positions or some other salient features. ^45^ Some key steps of building the MSM and determining the MFPT are described below in brief.

### Featurization and dimensionality reduction

For each frame of an MD trajectory, we built a 49-dimensional inter-helical distance feature by computing, for every selected C*_α_* on one helix, the minimum C*_α_*–C*_α_* distance to the opposing helix. Distance data from all systems (apo and different Py-Pip-bound Nlns) were concatenated and used as features to train a time-lagged independent components (tICA) model using the Deeptime package to capture the slow collective motions in a shared space.^46–48^ The average of these 49 minimum distances is defined as the interhelical distance (*d_IH_*), which is also used as a collective variable for estimating the free energy in apo and Py-Pip bound Nln later in this work.

### State discretizations and transition probability matrix

The tICA space was then partitioned with *k*-means clustering using 50 ≤ *k* ≤ 500 microstates. Here, the clustering algorithm was trained on the combined, all-system dataset so that state definitions are consistent across systems. For each system separately, we built counts matrices **C**(*τ*) by counting the transitions between states along the MD trajectories at lag times (*τ*). MSMs were built using values of *τ* = 1, 5, 10 and 20 ns. The counts matrices were column-normalized to generate the respective transition probability matrices.

### Macrostate definition and MFPTs

The states (clusters) were labeled as *closed* if their average interhelical distance *d*_IH_ ≤ 1.15 nm and *open* if *d*_IH_ ≥ 1.55 nm. The set of *closed* states form the *closed* basin and set of *open* states form the *open* basin for any typical system. These macrostates are used to compute mean first passage times (MFPT) of open to closed and closed to open interconversion in the Results section. MFPTs were estimated using the *calc mfpt* function of our CSNAnalysis package.^49^

### Parameter choices and sensitivity

Unless noted otherwise, MSMs in the Results section use *k* = 200 microstates and *τ* = 1 ns. Varying *k* within 50–500 did not change the qualitative MFPT trends across systems (SI Fig. S1). In contrast, longer lag times resulted in highly inflated MFPTs, often by 10–50× the simulation timescale, indicating that coarse temporal sampling misses kinetically important, faster transitions (SI Fig. S2). We therefore have adopted *τ* = 1 ns to preserve transition information.

### Conformational Space Networks

The conformational space networks (CSN) are quantitative representations of transition probability matrices obtained using the CSNAnalysis package. ^49^ CSNs are used to visualize the observables from MSM such as microstate properties, weights of the states and the probability and rate of transition between states. In this study, we used CSNs to explain the kinetics of channel opening/closing through the relative stability of the *open* or *closed* basins and the rate of interconversion for different Py-Pip bound systems. CSNs are visualized with the Gephi software as 2-D graphs consisting of edges and nodes.^50^ Each node in the CSN represents a microstate of the Markov model and each edge represents the transition probability between two states (nodes). In this work, the size of the nodes are scaled to their steady state probability and the width of the edges shows the transition probability. The color scheme in the CSNs follows the interhelical distance parameter (*d_IH_*) as it directly relates to the *open* and *closed* conformations.

### Differential scanning fluorimetry (DSF)

Differential scanning fluorimetry was performed to understand the effect of activator binding on the melting temperature (T*_m_*) of Nln, following a modified protocol undertaken in a recent study by Jayaraman et al. ^25^. The SYPRO Protein Stain (Invitrogen, product S6651) is used in our DSF experiments. A total 20 µL of assay mixture containing 3 µM recombinant Nln in artificial cerebrospinal fluid aCSF (126 mM NaCl, 26 mM NaHCO_3_, 3 mM KCl, 1.4 mM KH_2_PO_4_, 25 mM HEPES, 4 mM glucose, 1.3 mM MgCl_2_, 1.4 mM CaCl_2_, 0.2 µM ZnSO_4_ at pH 7.2) with 0.01 % final assay concentration of Triton X-100, Py-Pip and 20X SYPRO Protein Stain were mixed in a 96-well PCR plate and read in CFX96 Touch*^T^ ^M^* Real-Time PCR Detection System (Bio-Rad). The reaction mixtures without Py-Pip served as control. The samples were thermally denatured by heating from 25℃ to 99℃ at a ramp rate of 0.55 *^◦^*C/min and SYBR/FAM filter setting of 470 ± 20 nm excitation and 520 ± 10 nm emission was used for readings at each heating step.

## Results

### Activator binding in Nln: Melting temperature, binding sites and free energies

We first aim to confirm that in the process of enhancing the catalytic activity of Nln, Py-Pip indeed binds to the protein. Hence we carried out DSF experiment with 10, 30 and 100 *µ*M of Py-Pip in Nln and also with control Nln without Py-Pip, replicating each experiment four times. Fig. 2A shows that the melting temperature (T*_m_*) of Nln shifts consistently as we increase Py-Pip concentration, with all testing concentrations showing statistically significant lowering of T*_m_* compared to the control. The melting temperature (T*_m_*) of the control Nln is 59.3 ± 0.1 *^◦^*C, which subsequently decreases to 58.4 ± 0.2 *^◦^*C, 56.7 ± 0.3 *^◦^*C, and 54.6 ± 0.8 *^◦^*C in presence of 10, 30, and 100 *µ*M of Py-Pip, respectively.

**Figure 2:**
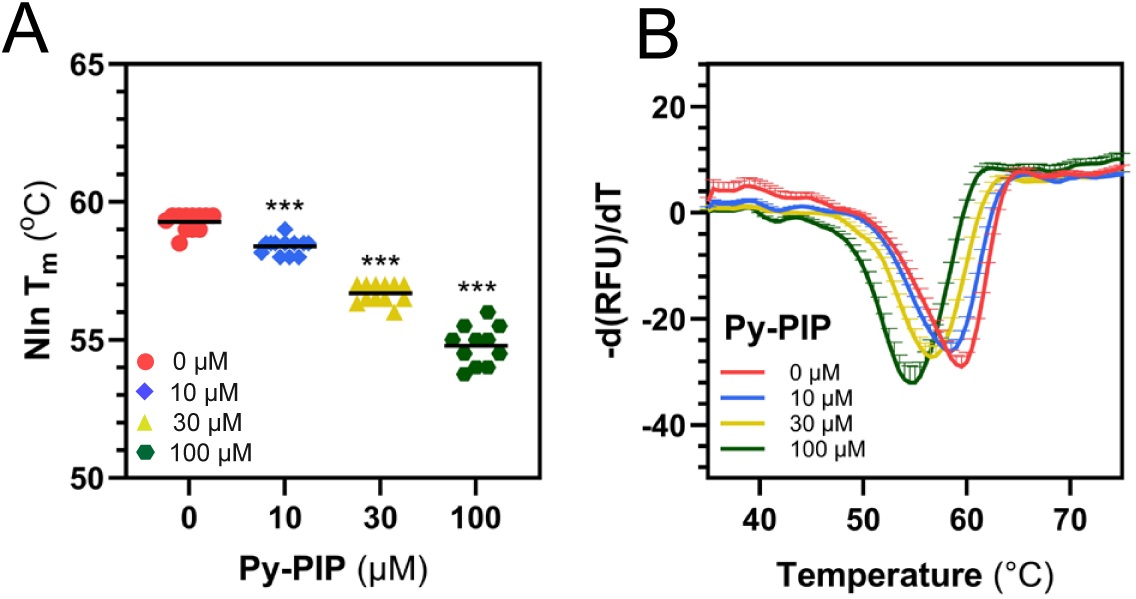
DSF analysis of Py-Pip and Nln interaction: (A) Concentration-dependent effect of Py-Pip (at 10, 30 and 100 *µ*M) on the melting temperature (T*_m_*) of recombinant Nln across 4 independent experiments. The three asterisks indicate a p-value *<* 0.001 compared to the control condition. (B) Negative derivative (-d(RFU)/dT) curves at corresponding Py-Pip concentrations, where RFU is the relative fluorescence unit.

The substantial decrease (∼5 *^◦^*C at 100 *µ*M Py-Pip) in the T*_m_* of Nln indicates direct Py-Pip binding, here resulting in thermal destabilization of Nln. The first derivative of relative fluorescence units (Fig. 2B) shows a smooth transition in all four cases, with the peak minima shifting consistently towards lower temperature at higher Py-Pip concentration. Hence, the thermal unfolding profiles of control Nln and Nln in three different Py-Pip concentrations clearly indicate that Py-Pip binds to Nln, and also reduces the stability of the protein in a concentration-dependent manner.

Next, we investigated possible binding sites of Py-Pip in Nln by running 10 independent replicates of 1.2 *µ*s long unbiased MD simulations, where 10 Py-Pip molecules are added randomly in the water-box containing Nln, leading to a ligand concentration of 0.022 M. These unbiased simulations revealed specific Nln regions that are preferred for binding by the activator molecules. Fig. 3A shows the probability densities of the activator utilizing the VolMap tool of Visual Molecular Dynamics (VMD) software with a cutoff of 0.025.^51^ In this figure, the green voxels within the contours have a *>* 2.5% probability of containing an activator atom across all simulations. From this analysis, we identify three most probable binding regions of Py-Pip in Nln that are consistent across all the runs and denote them as sites I, II and III. Site I has no direct interaction with the interhelical or catalytic residues, but shows a large, high-occupancy binding region. Site II has interactions with the residues in the inner core of the interhelical region and is closest to the catalytic site of Nln. However, it does not interact with any residues in the vicinity of the catalytic site. Site III has interactions with the residues in the interhelical region on the exterior side (upper right end). In Fig. 3B, we quantify the significant intermolecular interactions between Nln residues and the activator molecules across all 10 runs. The set of significant residues is determined by the threshold that the residue and Py-Pip contact must be present in at least 3% of cumulative simulation data, and a contact is defined when the Nln:Py-Pip heavy atom distances are less than 3.5 Å.

**Figure 3:**
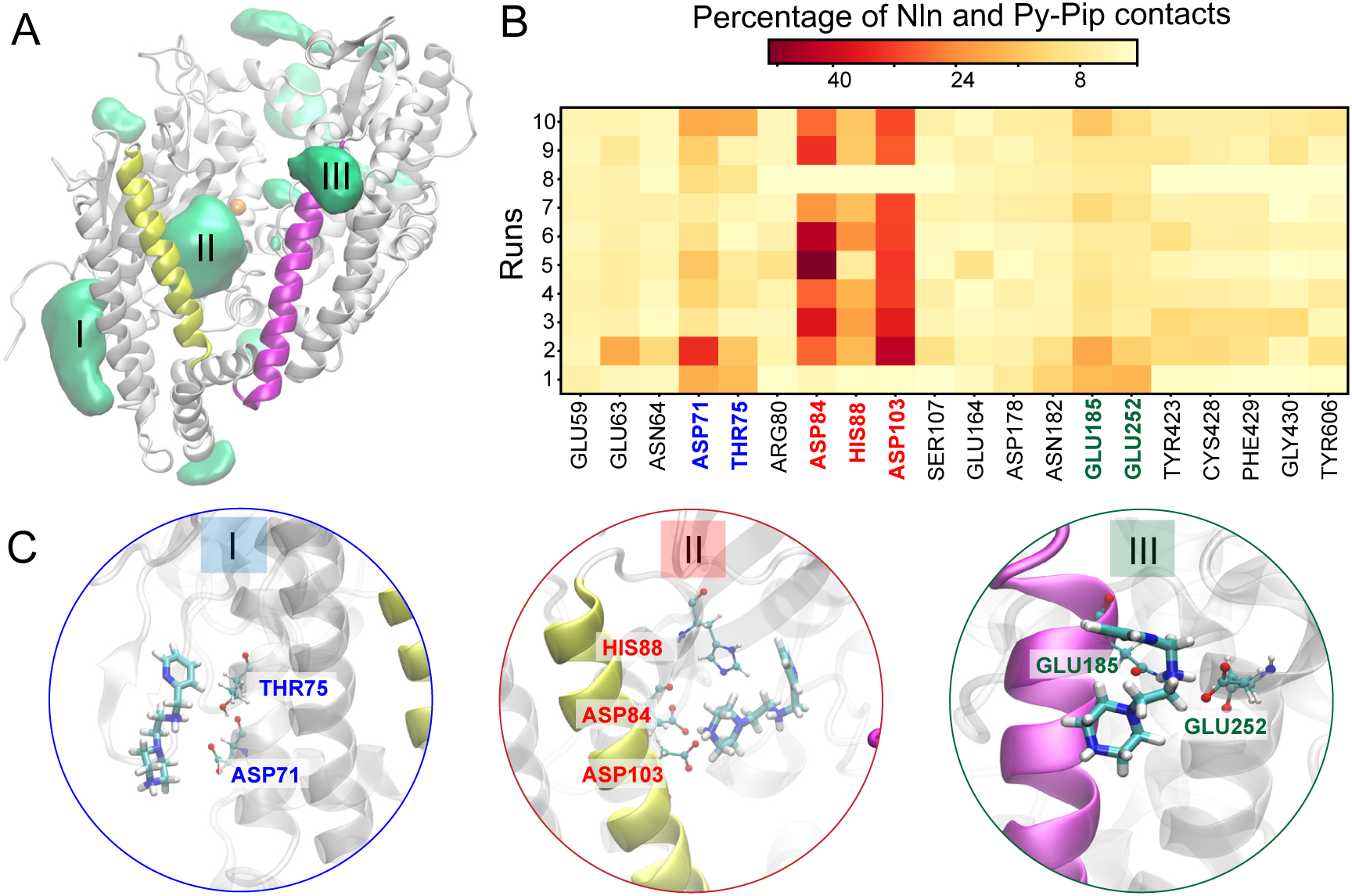
Py-Pip bound Nln: (A) Probable binding regions of Py-Pip activators in Nln are shown in green with the numbers (I, II and III) as identifiers, (B) Nln Py-Pip intermolecular interaction map for each of the Nln:10Py-Pip simulations (C) Representation of most probable Nln and Py-Pip interactions in binding regions I, II and III as seen in (B).

As the Py-Pip molecule has both H-bond donor and acceptor moieties, the majority of the significant binding site residues of Nln are polar or charged at physiological pH. Importantly, the most prominent interacting residues (e.g. ASP71, ASP84, HIS88, ASP103, and GLU185) are conserved across 8 out of 10 runs. This implies the relevance of these residues as the major binding sites of Py-Pip in Nln. The most probable Py-Pip orientations in each of the sites are shown in Fig. 3C.

In order to investigate how the binding of Py-Pip affects the overall Nln conformation space and dynamics, we first compare root mean squared fluctuation (RMSF) curves for apo Nln and Nln:10Py-Pip simulations (Fig. 4A). Throughout the protein, we observe higher RMSF values in the Nln:10Py-Pip simulations, indicating that the average effect of Py-Pip at high concentration is to make Nln more dynamic compared to apo. This is particularly pronounced in the interhelical channel region (residues 20-230, shown in black in the inset), which is known to play a major role in Nln opening and closing motions. There are also two flexible loops that have significantly higher RMSF in the Nln:10Py-Pip ensemble, shown in the inset (right) of Fig. 4A. Interestingly, both of these loops are adjacent to the two binding sites (I and III) of Py-Pip that may play a role in enhancing loop motions. It should be noted that the channel region interhelical residues, which form the multiple salt-bridges and drive the interconversion of *open* and *closed* ensembles, show some of the highest deviation in the RMSF in the presence of activators from the apo ensemble. Hence, we have used these residues to construct the interhelical distance (*d_IH_*) feature to characterize the *open* and *closed* states, and the transition between them under different activator-bound conditions. The detail of computing *d_IH_* is discussed in the Methods section. Interestingly, in the presence of the Py-Pip activators, the free energy along *d_IH_* changes significantly compared to the apo Nln. As seen in Fig. 4B, while the *closed* conformations of Nln are favored in the apo ensemble (red), in the Py-Pip activator-bound Nln ensemble (blue), the open conformations have significantly lower free energy. The preferential stability of *open* states points towards a probable mechanism of activation by Py-Pip, where the substrate entry into in the catalytic site could be facilitated by activators that favor an open channel configuration. The overpopulation of *open* states in Nln:10Py-Pip ensemble is in accordance with the previously observed lowering of Nln melting temperature during DSF experiments in presence of Py-Pip.

**Figure 4:**
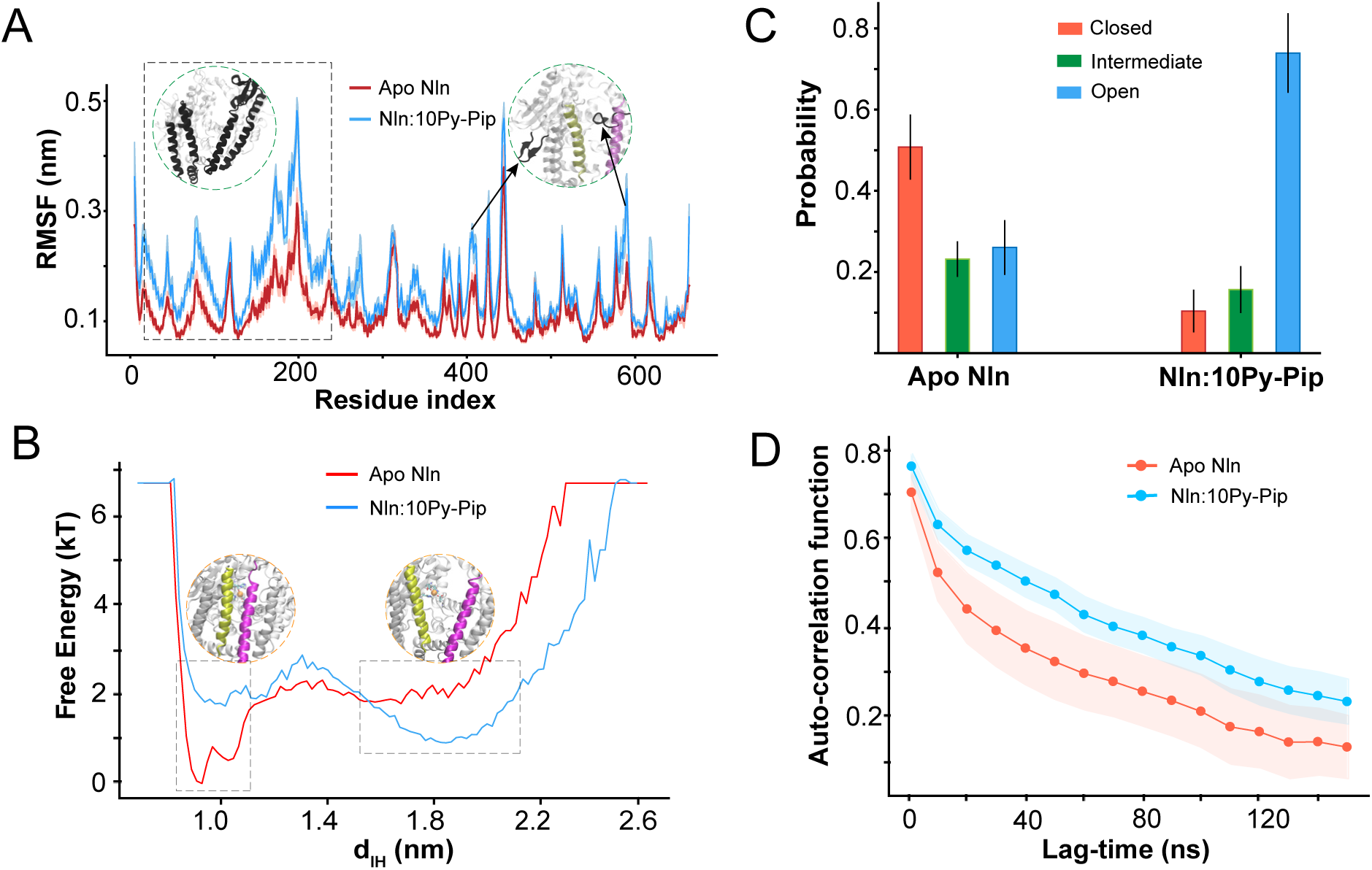
Comparison of Apo and Nln:10Py-Pip simulations. (A) Root mean square fluctuation (RMSF) in Nln residues in apo Nln simulations (red) and Nln:10Py-Pip simulations (blue). The shaded regions indicate error bars across runs. The regions of larger difference in RMSF are shown in black in the 3D protein structures (inset). (B) The free energy land-scape of apo Nln (red) and Nln:10Py-Pip simulations (blue) along the interhelical distance collective variable. *Open* and *Closed* basins are shown in inset. (C) Probability distribution of *Open*, *Intermediate* and *Closed* states shown in red, green and blue colors for apo Nln and Nln:10Py-Pip simulations, with the error bars showing the standard errors of the mean across runs. (D) The auto-correlation of the backbone atoms in the interhelical region calculated from apo Nln (red) and Nln:10Py-Pip (blue) simulations.

To better understand the conformational impact of Py-Pip binding, we quantify the fraction of the *open* and *closed* conformations in the apo and Py-Pip bound Nln ensembles. As mentioned in the Methods section, we define Nln conformations with *d_IH_* ≤ 1.15 nm as part of the *closed* basin, while the *open* basin is defined by *d_IH_* ≥ 1.55 nm. Any conformation in the range 1.15 *< d_IH_ <* 1.55 nm is defined as *intermediate*. These cut-offs are obtained from the *open* and *closed* basins in the free energy plots of apo and Py-Pip bound Nln in Fig. 4B. Subsequently, we plot the average probability of the *closed*, *intermediate* and *open* states across all the runs in the apo and Nln:Py-Pip simulations in Fig. 4C. As expected from the free energy landscape, the *closed* state of Nln in apo simulations is favored (with 50% probability, compared to 25% each in the *open* and *intermediate*) whereas in the presence of Py-Pip molecules, Nln has a *>* 70% probability of adopting an *open* conformation.

In order to examine the effect of activator on Nln kinetics, we calculate auto-correlation functions using the positions of C*_α_* atoms in the interhelical region. Interestingly, we find consistently higher auto-correlation across lag-times in the presence of Py-Pip (Fig. 4D). This shows that despite higher fluctuations (RMSF), the Nln:10Py-Pip system overall demonstrates a longer memory of conformations, with kinetically slower opening and closing motions. This highlights that the open state in the presence of Py-Pip is not necessarily more dynamic and can be thought of as a relatively stable conformational basin, in accordance to the observed free energy landscape of Nln:10Py-Pip (Fig. 4B).

### Binding-site-specific effects on Nln conformational fluctuations

Although the Nln:10Py-Pip simulations above were well-suited to identify putative binding sites, many frames involved multiple Nln and Py-Pip interactions and it is difficult to assess which specific activator molecules are most responsible for the observed Nln conformational changes. To understand the effect of specific Py-Pip binding we set up three Py-Pip-bound Nln systems, where a single Py-Pip molecule occupies one of the three binding regions shown in Fig. 3A. To avoid ambiguity due to different starting conformations of the three systems, we carefully chose one snapshot (Fig. 5A) from the original simulation with 10 copies of Py-Pip (Nln:10Py-Pip) simulation, where all three regions are simultaneously occupied by three different Py-Pip molecules. Each specific Py-Pip-bound Nln system is prepared by retaining only a single Py-Pip molecule. Following the nomenclature in Fig. 3A, we name the three systems as system I, II and III. Since Nln adopts an open conformation in this starting point, which is significantly different from the apo simulation starting point with more than 10 Å RMSD of the backbone of channel region residues, we also conducted additional apo Nln simulations from this open conformation to serve as a reference. Henceforth, we denote this system as reference Nln or Ref Nln. For each of the four systems (reference Nln and the 3 Py-Pip-bound Nln systems), we carried out 20 runs with 600 - 900 ns of simulation time each. To ensure that we observe the effect of Py-Pip bound at the specified region, we exert a flat-bottom potential on the Py-Pip atoms that restrains the molecule within a sphere of 7.5 Å from specific Nln residues in the binding site. The other details of the simulation setup such as temperature, pressure, box size, ionic strength are the same as those provided in the Methods section.

**Figure 5:**
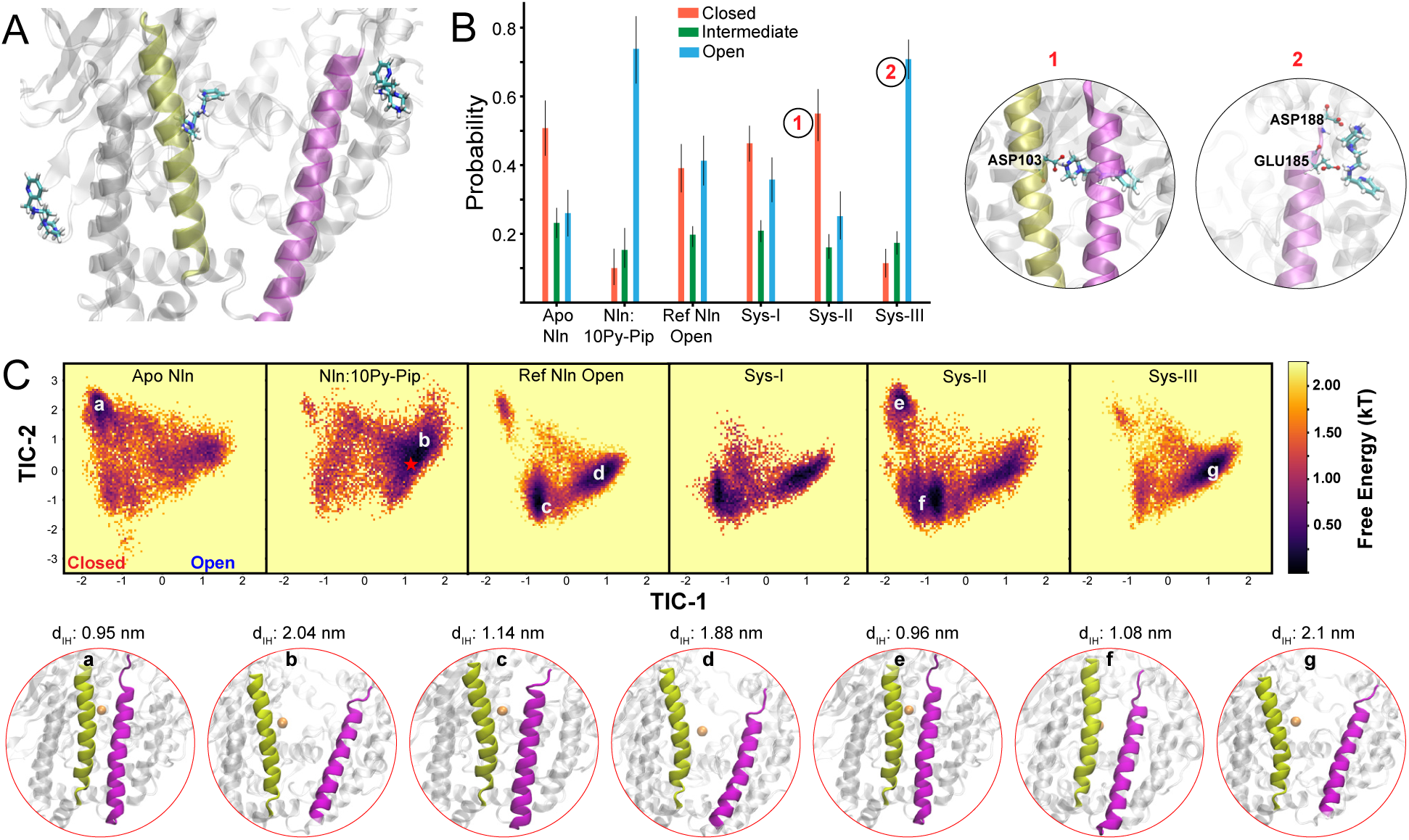
(A) The reference starting conformation, where all three binding regions are simultaneously occupied. (B) The probability distribution of *Open*, *Intermediate* and *Closed* states shown in red, green and blue, respectively. These are shown for apo (crystal), apo (open), System I, System II, System III and Nln:10Py-Pip. The most probable poses in the closed System-II and open System-III are shown in the adjacent circles. (C) Free energy distributions along TIC-1 and TIC-2 for all six systems in our study with representative Nln conformations in all the relevant basins shown in the lower panel.

We quantify the fraction of the *open* and *closed* conformations in the 3 systems, along with the reference Nln ensemble in Fig. 5B. While system I, where the Py-Pip is bound in the outer region, shows no major deviation in the ratio of *open* and *closed* conformations compared to that of the apo reference simulation, systems II and III produce significant differences in the fractions of *open* and *closed* conformations. The trend in system III is identical to that of the Nln:10Py-Pip ensemble, where we observe a significantly favored open conformation with more than 75% probability. In system II, where the Py-Pip is bound to the inner core of the interhelical channel, we notice that the probability of the closed conformations increase significantly. This is an intriguing observation as it indicates a possibility of an alternate mechanism of activation of Nln by the Py-Pip molecules. It has been previously noted that the closed conformation is essential for the catalytic reaction of substrates inside the Nln cavity.^6,9,13^ Hence, while the Nln:10Py-Pip and system III ensembles indicate Py-Pip has a propensity to keep the channel open, possibly allowing more facile substrate entry and/or product release, the system II ensembles reveal that Py-Pip can also catalyze the *open* to *closed* transition, which could also enhance the rate of the catalytic reaction, after binding of the substrate. The most probable Py-Pip binding poses in system II and III are shown in the right hand panel of Fig. 5B. In the closed conformation of system II, Py-Pip is most likely to form H-bonds with the ASP103 residue of the inner side of the left helix. In the open conformation of system III, Py-Pip can form multiple H-bonds with GLU185 or ASP188 residues in the top left corner of the right helix.

Furthermore, we carry out a combined time-lagged independent component analysis (tICA) with all six different ensembles to understand the landscape of Nln conformations of each of the systems. A 5-dimensional tICA model is trained using the combined dataset of interhelical distance features (49 per frame) to identify the slowest collective modes. We observe that the first component (TIC-1) separates out the *open* and *closed* conformations (SI Fig. S3). The second component (TIC-2) describes an out-of-plane angular shift of the two helices (SI Fig. S4). Free energies in the TIC-1 and TIC-2 space are plotted in Fig. 5C for the six different systems, with representative conformations from the high-probability basins shown in the lower panel along with their respective interhelical distances (*d_IH_*). As expected, the apo Nln has a stable *closed* basin (a) and also samples the broad open region fairly well. On the other hand, the Nln:10Py-Pip ensemble primarily samples the shallow *open* basin (b). The reference apo Nln ensemble starting from an open conformation samples majorly a *semi-closed* basin (c) and the *open* (d) basin. This *semi-closed* basin (c) has minor structural differences from the *closed* basin (a) with an out-of-plane angular shift between the two helices marginally increasing the interhelical distance, thus very slightly opening the channel even in the *closed* state, with *d_IH_* ≤ 1.15 nm. Hence, we denote this basin as *semi-closed*. System I has almost an identical sampling pattern as the reference Nln ensemble and system III primarily samples the shallow *open* conformations (g). Interestingly, system II has significantly stable basins at both *closed* (e) and *semi-closed* (f) conformations, with reasonable sampling of the *open* states. Remarkably, the other Py-Pip bound systems along with the reference Nln ensemble have a minimal sampling of the primary, *closed* basin (a or e). Moreover, the Py-Pip bound Nln simulations started from an extreme open state as highlighted by the red asterisk in the Nln:10Py-Pip free energy plot. This suggests that Py-Pip binding to the inner core (system II) of the interhelical channel could lead to *closed* conformations, similar to apo Nln, that could potentially reorient the substrate to a favorable conformation within the cavity and enhance the Nln catalytic activity.

### Kinetics of activator-mediated conformational changes

The above results suggest a complex interplay between multiple processes occurring on similar timescales: activator (un)binding, Nln conformational change, substrate binding, catalysis and product release. In such a scenario, the rate of interconversion between the *closed* and the *open* state becomes an important quantity for modeling the action of the activators in different binding sites. One can quantify the rate of *closed* → *open* and *open* → *closed* transitions by calculating the first passage times (FPT) of the respective transitions along a continuous trajectory. Averaging over all such FPTs will provide us with the mean first passage time (MFPT), which we can compare between our simulation ensembles. However, the interconversion of the *closed* and *open* states is slow compared to the simulation timescale and one can only observe a handful of these transitions even with microseconds of molecular simulation. Hence, estimating FPTs solely based on counting transitions seen in the trajectory leads to statistically noisy mean first passage times (MFPT) with high standard errors (SI Table S2).

Markov state models provide a robust estimation of the underlying slow kinetic processes even when the transitions are observed rarely in the simulation trajectories.^45,52^ The details of constructing Markov models using the MD simulation data is described in the Methods section. In Fig. 6A, we show the MFPTs of *closed* → *open* and *open* → *closed* transitions in the Py-Pip-bound ensembles and the corresponding apo reference ensemble. Following the definitions from Methods section, a *closed* state has *d_IH_* ≤ 1.15 nm and an *open* state has *d_IH_* ≥ 1.55 nm. We notice that system II has a 2-fold increase in the MFPT of the *closed* → *open* transition and system III lowers the same by 4-fold compared to reference Nln. This indicates that while Py-Pip binding inside the channel can slow down the channel opening, Py-Pip binding at the upper end of the channel can enhance the opening process significantly.

**Figure 6:**
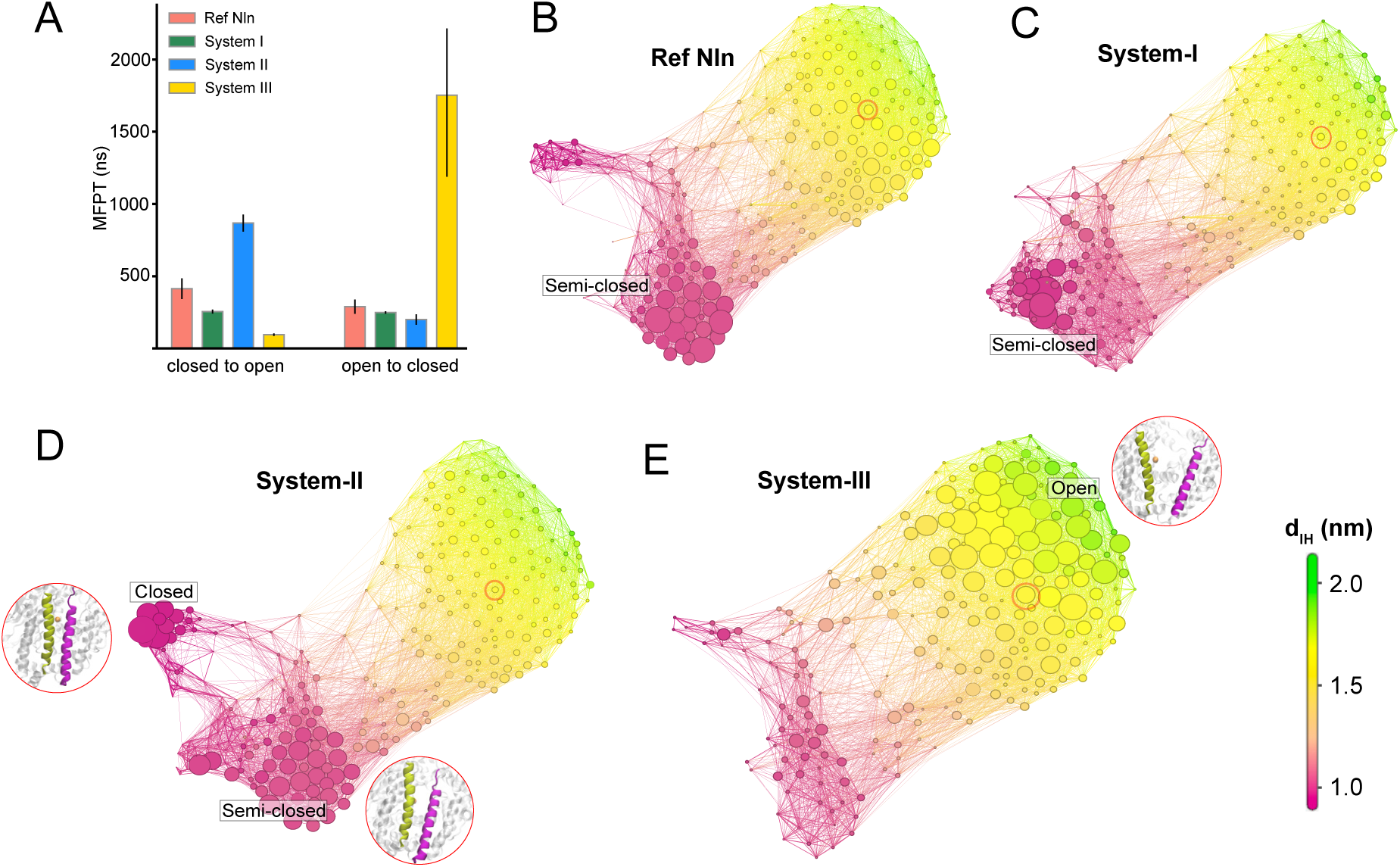
(A) MFPT estimated from the Markov model for *closed* → *open* and *open* →*closed* transitions. The error-bars indicate standard errors observed across a set of independent runs for each of the systems. (B) - (E) Conformational space network (CSN) obtained from the individual transition probability matrices of reference Nln, system I, system II and system III ensembles respectively. The localized *closed*, *semi-closed* and delocalized *open* basins are pointed in adjacent boxes. Representative snapshots of system II and III are provided in the inset. The red circled state contains the same starting conformation of the simulations. The color scheme of the CSNs (interhelical distance parameter) shown in the bottom right.

System I shows a slight decrease in MFPT of the *closed* → *open* transition. On the other hand, the kinetics of *open* → *closed* transitions are significantly altered in system III, which has a ∼ 10 times higher MFPT compared to the apo reference simulation. This implies that Py-Pip binding at the upper end (site/system III) of the interhelical channel keeps the channel open for longer time, which may have a favorable impact on enzyme activity by allowing the substrate to bind or the products to release. System I does not show much difference from the apo reference and system II lowers the MFPT of the *open* → *closed* transition slightly.

The kinetics of channel opening and closing can be explained by the relative stability of the *open* or *closed* basins at different Py-Pip-bound systems using the conformational space networks (CSN) as shown in Fig. 6B-E. The details of CSN construction and representation is discussed in the Methods section. It should be noted that the state discretizations are performed using a combined dataset with equal contributions from all six systems in our study. This ensures consistent state definitions across all the systems, allowing the CSNs to be directly compared. The node size in each CSN corresponds to the steady probabilities of those states (nodes) from the system-specific transition probability matrices. The states (nodes) that are not visited by a system during its simulation are omitted from the corresponding CSN. The Py-Pip bound Nln simulations are initialized from the same conformation in the *open* basin, which is highlighted by a red circle in Figs 6B-E. The dense distribution of edges in the *open* region in all the CSNs suggests that the *open* states have a broad basin with easier transition between the states. Interestingly, we observe the *closed* and *semi-closed* basins are tightly packed with high weighted nodes indicating existence of sharp energy minima. While in the reference Nln, the *closed* basin has low weighted states, we do not observe them in system I or system III at all. System II however has substantial density of states in both the *closed* and *semi-closed* basins. Moreover, unlike apo reference Nln where there are no connection between the *closed* and *semi-closed* basins, in system II we see significant number of edges connecting these two basins. This indicates, unlike system I and III, system II promotes interconversion between the *closed* and *semi-closed* ensembles, slowing the eventual *closed* → *open* transitions significantly. On the other hand, system III has significantly high probability nodes at *open* basin that is in accordance with our previous analyses.

## Discussion

The consistent and statistically significant shift in the melting temperature of Nln at different Py-Pip concentrations indicates a direct binding of Py-Pip to the protein. Our simulations reveal three discrete, plausible binding sites for the small-molecule activator on Nln, each involving distinct polar or charged residues and presenting unique functional relationships to the dynamics of the Nln interhelical channel. Fig. 5 suggests that site I, located in an exterior cavity near the N-terminus, appears to be conformationally neutral, leaving the open/closed equilibrium largely unperturbed. In contrast, sites II and III exert opposite effects on the global conformational landscape of Nln. Py-Pip binding at site II, inside the interhelical channel, stabilizes the *closed* conformation, while site III binding, located on the exterior side of the interhelical region, strongly favors the *open* conformations. Although Fig. 3B points to a relatively higher binding affinity of site II compared to site III, it should be noted that *open* conformations are required for activators to access site II, making it reliant on the *closed* to *open* interconversion. Site III, on the other hand, is accessible in both *open* and *closed* conformations. Hence, we suggest both binding sites should be considered in exploring the mechanism of action of Nln activators.

The existence of such opposing effects from different binding locations suggests a possible mechanism in which small molecules can modulate distinct sub-steps of the Nln catalytic cycle. The catalytic reaction in metallopeptidases proceeds through multiple coordinated events: *substrate entry* into the cleft, precise *substrate positioning* within the active site in the *closed* state, the overall *hydrolysis reaction*, and finally the *release of the cleaved peptide products*. Activators binding in site III could facilitate the *substrate entry* and/or *product release* step(s) by reducing the free-energy barrier for channel opening (Fig. 5). This shift in the free-energy landscape is in excellent agreement with the DSF experiments, where the destabilization of Nln by Py-Pip can be connected to the overpopulation of the *open* conformations. Moreover, as seen in Fig. 6A, Py-Pip binding at site-III also kinetically favors the *open* state, slowing down the *open* to *closed* transition by an order of magnitude compared to other Py-Pip-bound systems and apo Nln. On the other hand, activators binding at site II may enhance the subsequent catalytic reaction by favoring the *closed* states (Fig. 5B, C), slowing down the *closed* to *open* transition (Fig. 6A). Hence, site-II binding could allow the substrate to reorient in a chemically favorable pose against the active site residues and water molecules for bond scission. Significantly, the high-concentration Py-Pip simulations, in which both Site-II and Site-III binding occur, were found to stabilize the open state, in qualitative agreement with the DSF results.

The combined effect of activators binding in different pockets, with different effects on the Nln conformational equilibrium could potentially be modeled quantitatively using a reaction network model. In this model, a comprehensive set of states describing Nln conformation (*open* vs. *closed*), activator binding (apo, site-II, site-III), substrate binding (apo, bound) and catalytic reaction state (reactant, product) could be built, with a set of rates describing their interconversion. While precise definitions of all the kinetic constants would be a challenging task, estimates could be used to reveal qualitative mechanisms of activation, and to test basic hypotheses of the proposed dual-site activation mechanism. For instance: what binding affinity would be needed in site-III to have a substantial impact on the overall catalytic rate? It is worth noting that the catalytic step should likely be treated as irreversible, which drives the network out of equilibrium, necessitating a rate-based approach. This will be the subject of future work.

Our Markov state model analysis provides a foundation for this work, as we have determined how site-specific activator binding can affect the rates of Nln conformational transitions. Site III significantly slows down the *open* to *closed* transitions and also increases the rate of *closed* to *open* transitions, kinetically biasing the channel toward prolonged open states (Fig. 6). Site II produces an opposite signature, delaying opening and modestly accelerating closure of the channel. In the context of multi-site occupancy of the activators, simultaneous engagement of sites II and III could, in principle, couple rapid substrate recognition with efficient catalytic closure of the channel, a probable scenario recently discussed for other mechanosensitive enzymes with hinge motions like Nln. ^38^

Experimental attempts thus far have not yet been successful in crystallizing the Nln:Py-Pip complex as the activator drives the enzyme into a highly dynamic ensemble. Differential scanning fluorimetry experiments in this work have shown a consistent lowering of the Nln melting temperature with increase in activator concentration. This indicates an activator-induced reduction in the global thermodynamic stability of Nln, which is also reflected in the free energy landscapes of our simulations. Py-Pip occupancy at site III prolongs *open* states, breaking the stable inter-domain salt-bridge network, while site II biases the protein toward an alternative *closed* basin. The resulting conformational heterogeneity together with the modest, overall micromolar affinity of Py-Pip, which is the sum of all the site-specific affinities acting in a counter-cooperative fashion, probably prevents the formation of a single, well-ordered lattice competent for X-ray diffraction. Thus, while multi-site binding of Py-Pip may be instrumental to lift catalytic turnover in solution, it simultaneously frustrates Nln:Py-Pip co-crystallization by amplifying structural plasticity. Interestingly, the substrate-bound structures of Nln were crystallized recently, and provides an important structural reference for the *closed*, catalytically competent state.^13^ This suggests that the site II binding may stabilize similar geometries even in the absence of substrate, potentially pre-organizing the active site for the incoming substrates.

Together these results provide a mechanistic framework for rational design of next-generation activators of Nln. Future optimization of molecular features may be directed to either selectively target one of the sites, or designing bi-functional ligands that engage both regions, yielding synergistic effects on overall catalytic efficiency. A reaction network model proposed above would provide further knowledge towards focusing one or both sites. Moreover, modulating the activity of a number of metallopeptidases such as thimet oligopeptidase (TOP), angiotensin converting enzyme (ACE/ACE2) could offer pharmacological benefits in various disease contexts. These enzymes have structural similarity in making salt-bridges across the two domains enabling them to attain both *open* and *closed* conformations. Hence, under-standing the mechanism of activation by Py-Pip molecules can provide us a template for a systematic, rational development of small molecule modulators for not only Nln but also any structurally similar metallopeptidase. This multi-site, conformation-driven activation paradigm offers new opportunities to enhance the activities of the metallopeptidases, enabling drug-discovery efforts aimed at reducing the toxic accumulation of peptides in the human body.

## Acknowledgments

AD and SB acknowledge support from grants R01GM130794 from the National Institutes of Health. VTK and BJO are supported by the National Institute of Neurological Disorders and Stroke under award number R01NS106879. The Py-Pip compound (NSC 66183) used in this study was obtained from the National Cancer Institute (NCI) Division of Cancer Treatment and Diagnosis, Developmental Therapeutics Program (http://dtp.cancer.gov). The content is solely the responsibility of the authors and does not necessarily represent the official views of the National Institutes of Health.

## Notes

### Competing Interest Statement

The authors have declared no competing interest.

