## Supplemental Info for "Conformation Driven Enhancement of Neurolysin Activity in Presence of a Small Molecule Activator"

December 12, 2025

### 1. MFPTs estimated from MSMs built with variable numbers of states

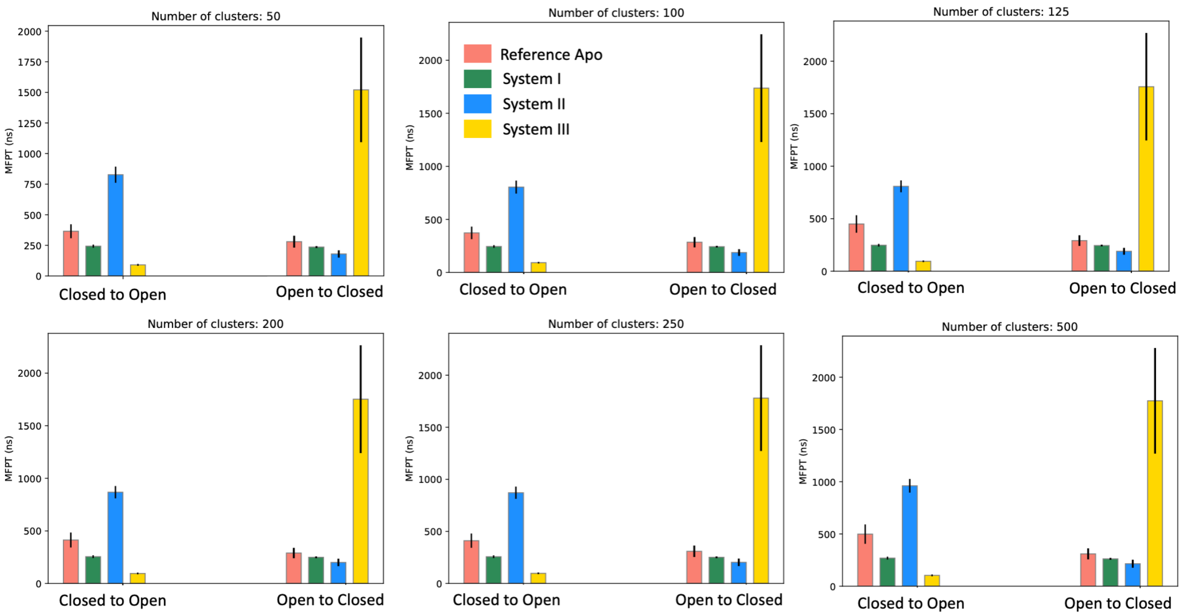


Figure S1: MFPT estimated from the Markov model for closed to open and open to closed transitions using numbers of clusters ranging from 50 to 500. The number of states has a minimal effect on the MFPTs.

#### 2. MFPTs estimated from MSMs built with different lag-times

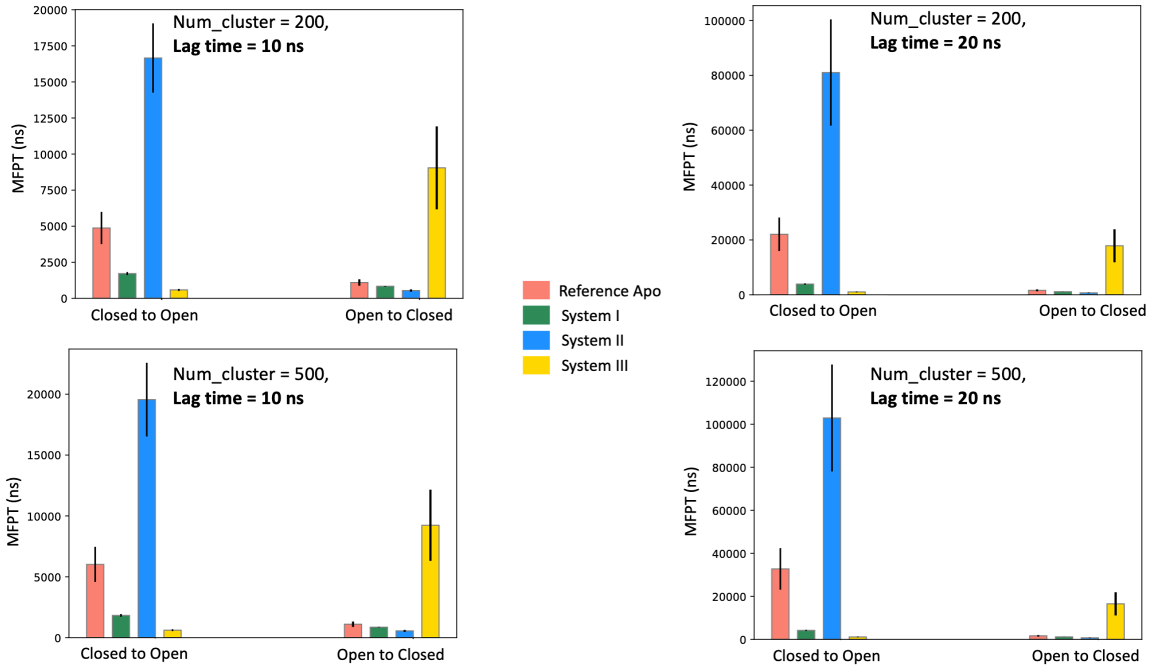


Figure S2: MFPT estimated from the Markov model for closed to open and open to closed transitions using different lag-times (10 ns and 20 ns). Longer lag times inflated MFPTs spuriously, often by 10–100× (e.g. sys-II), indicating that coarse temporal sampling misses kinetically important, faster transitions. This can happen as the open conformations construct a broad and shallow minima, where Nln can switch between 'semi-open' and 'open' states frequently. Hence, at longer lag-times, the model may end up missing the closed to open transitions.

#### 3. The TIC-0 variable describes Nln opening and closing

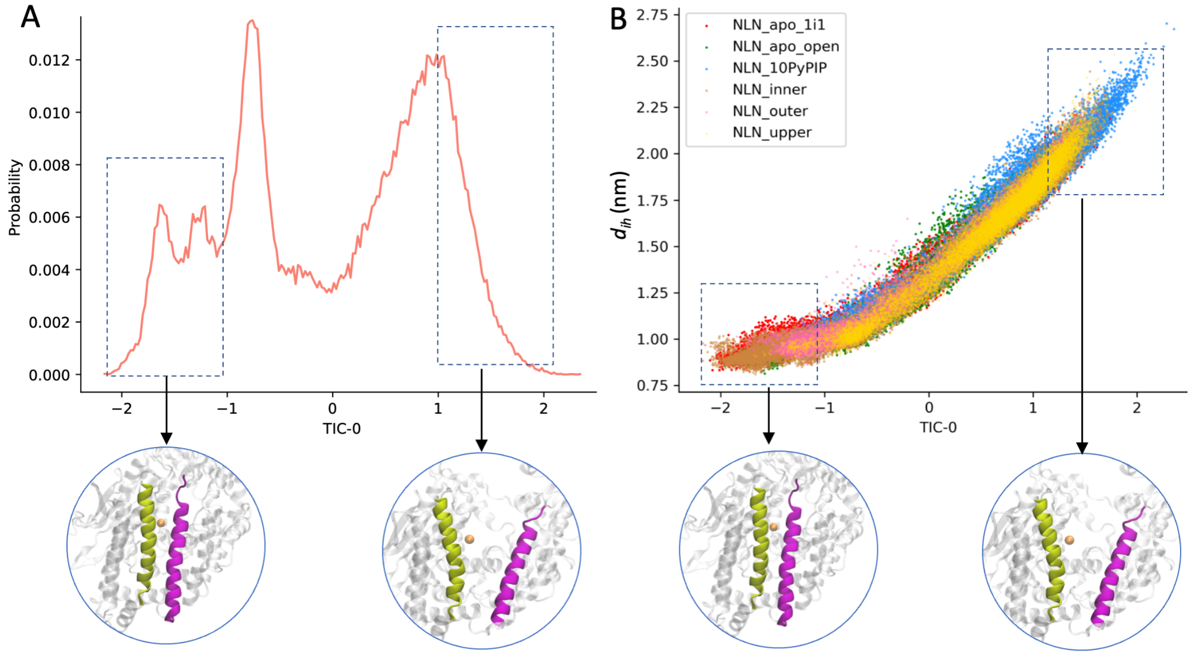


Figure S3: (A) A combined probability distribution across all systems of TIC-0 with two extreme ends of the distribution isolating closed and open Nln conformations. (B) Scatter plot of TIC-0 against $d_{IH}$ for all frames, individually shown for each of the six systems. This reveals a strong correlation between open and closed conformations with high and low value of TIC-0 respectively.

#### 4. TIC-1 describes an out-of-plane rotation of the channel helices

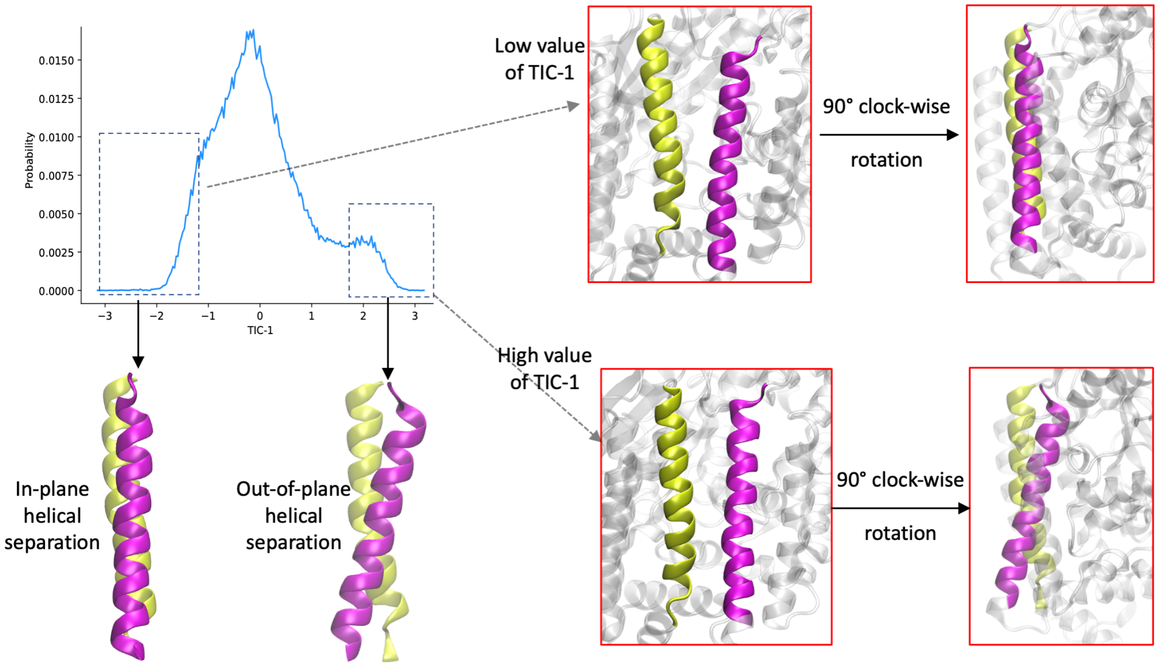


Figure S4: Probability distribution of TIC-1 with two extreme ends of the distribution describing an out-of-plane rotation in the channel region, shown by the only helical region residues below. Low values of TIC-1 (upper panel at the right) correspond to conformations where the two helices are parallel, while higher values (lower panel at the right) correspond to Nln conformations with rotated helices.

#### 5. Table of MD simulation detail

Table S1: Details of MD simulations with different systems

| System | Number of independent runs | Length of each run (ns) | Total simulation time ($\mu$s) | Concentration of the activator (M) |
|---|---|---|---|---|
| Apo Nln | 20 | 900 | 18 | N.A. |
| Nln:10Py-Pip | 10 | 1250 | 12.5 | 0.022 |
| Ref Nln (open) | 20 | 600 - 900 | 13.5 | N.A. |
| System I | 20 | 600 - 900 | 13.5 | 0.0022 |
| System II | 20 | 600 - 900 | 15.5 | 0.0022 |
| System III | 20 | 600 - 900 | 15.5 | 0.0022 |

#### 6. Table of MFPTs and standard errors directly estimated from the MD trajectories

Table S2: Direct estimation of MFPTs and their uncertainties (standard error) from MD trajectories without using MSMs. The error is at least 10× higher than what we see in case of MFPTs from MSMs.

| System | MFPT (ns) Closed to Open | Standard error (ns) Closed to Open | MFPT (ns) Open to Closed | Standard error (ns) Open to Closed |
|---|---|---|---|---|
| Apo Nln | 37 | 11 | 65 | 15 |
| Ref Nln (open) | 126 | 51 | 204 | 49 |
| System I | 51 | 28 | 302 | 111 |
| System II | 115 | 53 | 128 | 43 |
| System III | 122 | 95 | 339 | 143 |
| Nln:10Py-Pip | 31 | 21 | 314 | 106 |
